# Stress related network activity in the intact adrenal medulla

**DOI:** 10.1101/2021.04.13.439677

**Authors:** Jose R. Lopez Ruiz, Stephen A. Ernst, Ronald W. Holz, Edward L. Stuenkel

## Abstract

The adrenal medulla has long been recognized as playing a critical role in mammalian homeostasis and the stress response. The adrenal medulla is populated by clustered chromaffin cells that secrete epinephrine or norepinephrine along with other peptides into the general bloodstream affecting multiple distant target organs. Although the sympatho-adrenal pathway has been heavily studied, detailed knowledge on the central control and in-situ spatiotemporal responsiveness remains poorly understood. For this work we implemented electrophysiological techniques originally developed to elucidate CNS circuitry to characterize the functional micro-architecture of the adrenal medulla. To achieve this, we continuously monitored the electrical activity inside the adrenal medulla in the living anesthetized rat under basal conditions and under physiological stress. Under basal conditions, chromaffin cells fired action potentials with frequencies between ∼0.2 and 4 Hz. Activity was exclusively driven by sympathetic inputs coming through the splanchnic nerve. Furthermore, chromaffin cells were organized into arrays of independent local networks in which cells fire in a specific order, with latencies from hundreds of microseconds to few milliseconds. Electrical stimulation of the splanchnic nerve evoked the exact same spatiotemporal firing patterns that occurred spontaneously. Induction of hypoglycemic stress by administration of insulin resulted in an increase in the activity of a subset of the chromaffin cell networks. In contrast, respiratory arrest induced by anesthesia overdose resulted in an increase in the activity of the entire adrenal medulla before cessation of all activity when the animal died. The results suggest the differential activation of specific networks inside the adrenal gland depending on the stressor. These results revealed a surprisingly complex electrical organization and circuitry of the adrenal medulla that likely reflects the dynamic nature of its neuroendocrine output during basal conditions and during different types of physiological stress. To our knowledge, these experiments are the first to use multi-electrode arrays *in vivo* to examine the electrical and functional architecture of any endocrine gland.

**Significance Statement:** Stress from extrinsic (environmental, psychological) and intrinsic (biological) challenges plays a critical role in disturbing the homeostatic balance. While the body’s responses to stress are designed to ameliorate these imbalances, prolonged and dysregulated stress often drives adverse health consequences in many chronic illnesses. The better understanding of the sympatho-adrenal stress response, will potentially impact and improve the treatment of several stress related illnesses. This work focusses on the study of the functional architecture of the adrenal medulla, a key component in neuronal stress response.

## Main Text

### Introduction

The adrenal medulla has long been recognized as playing a critical role in mammalian homeostasis and the stress response. Numerous projections from the central nervous system regulate the activity of the preganglionic neurons, which innervate individual adrenal medullary chromaffin cells, thereby regulating catecholamine and peptide secretion into the circulation. The current study focuses on chromaffin cell electrical activity and networks in living anesthetized rats.

Chromaffin cells have been extensively studied *in vitro*, both in cell culture and in tissue slices, to determine biochemical, cell biological and biophysical aspects of cell function. Studies in tissue culture have illuminated biochemical and physiological pathways that lead to exocytotic release of catecholamines and proteins. Electrophysiological studies of cells in adrenal medullary slices have revealed that cells in their natural tissue environment have faster release kinetics (Moser and Neher 1997) and have variable degrees of electrical coupling (Moser 1998, Martin, Mathieu et al. 2001, Colomer, Martin et al. 2012, Hill, Lee et al. 2012, Guerineau 2018). Fast-scan cyclic voltammetry and amperometry detect individual epinephrine and norepinephrine fusion events in culture chromaffin cells (Wightman, Jankowski et al. 1991, Chow, von Ruden et al. 1992) and in slices of mouse adrenal medulla (Petrovic, Walsh et al. 2010).

Numerous studies have investigated biochemical and physiological function of chromaffin cells in intact preparations. Indeed, there is a rich history of *in vivo* investigations of biochemical changes in adrenal medullary content upon secretion following intense stress and of *in vitro* studies of secretion of catecholamines and proteins from perfused adrenal medulla (for reviews see (Smith and Winkler 1972, Viveros 1976); also (Wakade and Wakade 1982)). These studies established exocytosis as the mechanism of secretion in neuroendocrine tissue. More recently, an *ex vivo* preparation demonstrated spatially- and frequency- dependent epinephrine and norepinephrine secretion in the rat adrenal gland that is relevant to the stress response (Wolf, Zarkua et al. 2016). An *in vivo* study measured extracellular potentials from chromaffin cells in response to splanchnic nerve stimulation and suggested a role of electrical coupling in enhancing catecholamine secretion (Desarmenien, Jourdan et al. 2013).

In the present study we extended the investigation of adrenal medullary function *in vivo.* We have utilized silicon multielectrode array probes to record action potentials extracellularly that are generated by chromaffin cells upon electrical or stress-induce splanchnic nerve stimulation in anesthetized rats. We adapted signal processing techniques, which have been developed for elucidating central nervous system neuronal activation patterns, that enabled detection of simultaneous activity from as many as 40 chromaffin cells. There was a surprising richness of responses with spatially overlapping cell networks displaying distinctive electrical and physiological responses. The results suggest that rather than all chromaffin cells responding identically, there is a significant amount of specificity and tuning of chromaffin responses to physiological stimulation.

### Results

The adrenal medulla is populated by electrically excitable chromaffin cells that are organized into visually identifiable clusters (**Figure 1A, B**). The cells are approximately spherical and are innervated by the sympathetic nervous system via the splanchnic nerve that emanates from the spinal cord. Neuronal tracing studies suggest multiple neuronal circuits originating from the brain stem and cerebral cortex that could influence chromaffin cell electrical activity via the splanchnic nerve (Strack, Sawyer et al. 1989, Dum, Levinthal et al. 2019) and, thereby, provide control of specific cells or cell clusters upon different physiological stressors.

**Figure 1.**
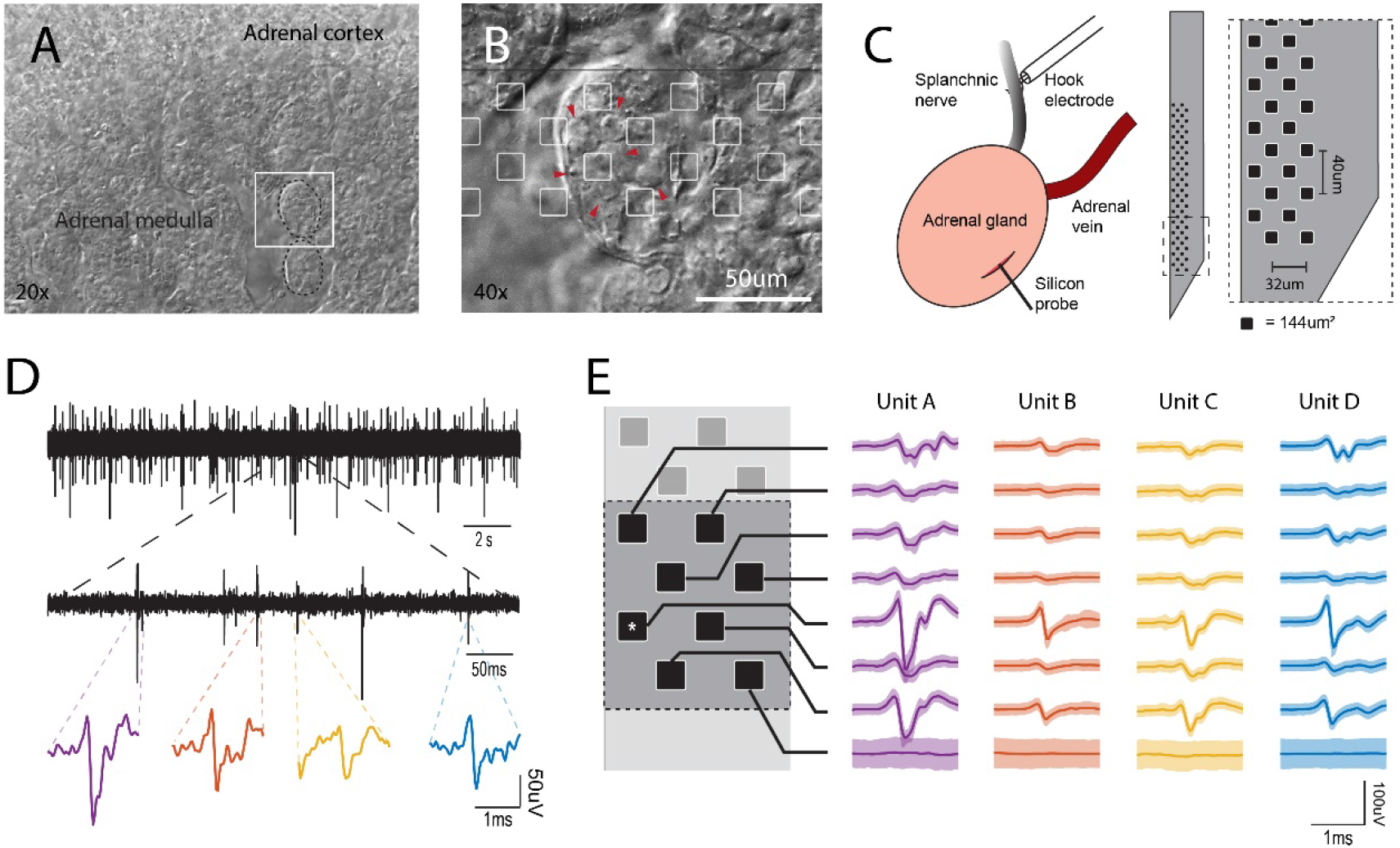
In vivo electrophysiological recordings from the intact adrenal medulla. A. Low power bright field micrograph showing the adrenal cortex and the adrenal medulla. The white square represents the area shown in panel B. B. Bright field micrograph showing the clustered architecture of the adrenal medulla (arrowhead = individual chromaffin cell). The silicon probe layout is shown in scale with the tissue, recording sites are presented as white squares. C. Left, preparation in the anesthetized rat for simultaneous stimulation to the splanchnic nerve and the recording of the electrical activity in the adrenal medulla. Right, 64 channel silicon probe layout, the area of each individual recording site is 144μm^2^. D. Spontaneous extracellular action potentials. Shown are recordings at low, medium and high expanded time scales. The most expanded recordings (bottom traces) show the waveforms of individual events. E. A cell’s action potential projects distinct waveforms depending on its position to each recording site in the neighborhood (the asterisk represents the channel shown in D), the set of these slightly different waveforms define a unit’s template and it is unique for each cell (solid line ± 2SD).

To investigate the electrical activity of chromaffin cells within the adrenal medulla of a living animal we positioned a 64-channel silicon multi-electrode array that was guided through the capsule and the adrenal cortex into the medulla (**Figure 1C**). The recorded signals displayed multiple abrupt downward voltage deflections or spikes with no evident firing pattern (**Figure 1D**). The size (generally 50 μV −100 μV) and speed (a millisecond or less) of the deflections indicate the signals reflect actions potentials from excitable cells (Gold, Henze et al. 2006). Due to the anatomy of the adrenal medulla, a single electrode often detected action potentials from multiple neighboring cells (**Figure 1B**). Conversely, an action potential from an individual chromaffin cell could usually be recorded in multiple channels, projecting different waveforms, as a result of passive temporal-spatial decay in the extracellular space, to each recording site (**Figure 1E**). This redundant information underlies template matching, the sorting technique used in this study to analyze responses of individual cells and their interactions (see Methods). During a single experiment, template matching enabled the detection of multiple cells, in some cases greater than 40.

#### Spontaneous electrical activity

Chromaffin cells fired in the absence of physiological stress (**Figure 2A**). The firing rates of individual cells varied widely, with ninety-five percent of the cells having firing rates between 0.2 and 4 Hz (550 out of 576), with a median of 1.36 Hz (**Figure 2B**). The baseline activity is consistent with the presence of circulating epinephrine even in the absence of stress.

**Figure 2.**
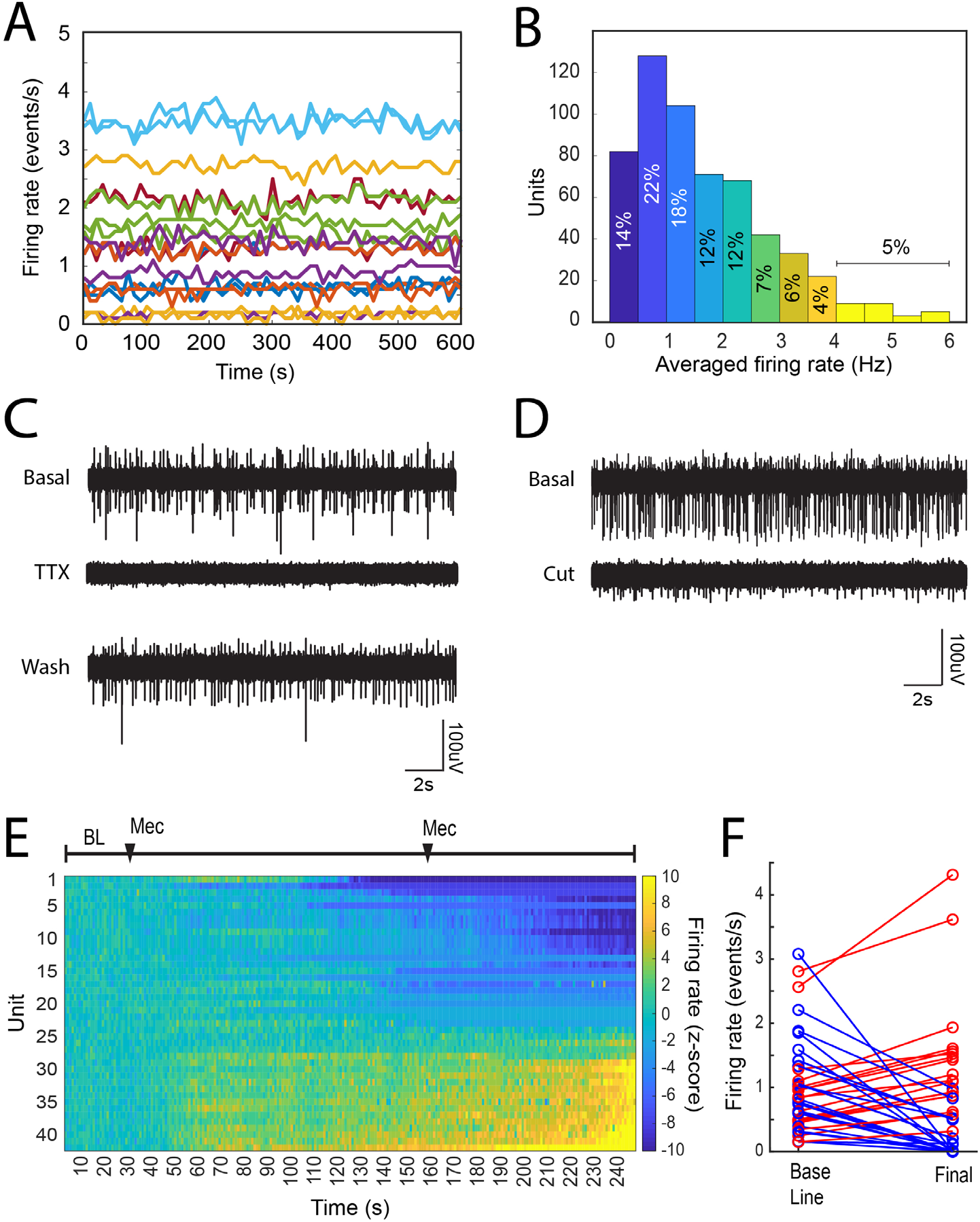
Basal chromaffin cell activity is driven by the splanchnic nerve. A. Firing rate in 10 s bins from a set of units recorded for 10 minutes under basal conditions in a single preparation. B. Averaged firing rate distribution from 576 units recorded during 40 preparations. C. The local application of the sodium channel blocker TTX, reversibly blocks the activity in the adrenal medulla. D. Severing the splanchnic nerve proximally to the adrenal gland, irreversibly abolishes the electrical activity in the adrenal medulla (timing). E. The systemic application of the nicotinic antagonist Mecamylamine hydrochloride inhibits a subset of the chromaffin cells population in the adrenal medulla. The firing rate was estimated in 10 s bins and the z-score (see Methods) was calculated by considering the first 5 minutes of the recording as baseline. After the baseline period two 2 mg intraperitoneal doses of Mecamylamine were administered at 5 and 25 minutes. F. Shows the averaged firing rate from each unit during the baseline period vs the final 5 minutes of the recording (blue=inhibited, red=increased).

We investigated the basis of the spontaneous activity. Application of the Na^+^ channel blocker tetrodotoxin (TTX) to the exposed nerve, reversibly inhibited the electrical activity, often reducing to zero the firing rate of individual chromaffin cells (**Figure 2C**). Activity was observed to partially recover following TTX removal and local washing of the splanchnic nerve. Cutting the splanchnic nerve caused a strong reduction of all spike activity. Remaining activity likely resulted from residual axonal activity and/or incomplete denervation based on anatomical branching of the splanchnic nerve (**Figure 2D**). We conclude that spontaneous activity in chromaffin cells is driven by basal levels of activation through the splanchnic nerve and not by autonomous activity of chromaffin cells.

A major excitatory neurotransmitter released from the splanchnic nerve terminals is acetylcholine, which interacts with the nicotinic cholinergic receptors on the plasma membrane of chromaffin cells. The systemic administration of the nicotinic receptor antagonist mecamylamine (Mec) reduced the firing rate of many but not all chromaffin cells (**Figure 2E and 2F**). Unexpectedly, the firing rate of some cells increased, suggesting a compensatory, non-nicotinic cholinergic pathway for chromaffin cell activation and/or sensitivity of specific central nervous afferent controls to mecamylamine. In either case this indicates specific and potentially differential control of networks of chromaffin cells in the adrenal medulla.

#### Network activity in the adrenal medulla

Complex waveforms were frequently recorded at individual electrodes **(Figure 3A)**. The templates derived from these recordings displayed spikes occurring in fast succession across a neighborhood of recording sites (*e.g.* orange and blue templates in **Figure 3B and** 3C). The sorting algorithm was able to resolve these complex waveforms into individual units (*e.g.*, cells) that fire in a specific sequence with very precise latencies (*e.g.* a1→a2; b1→b2→b3 in **Figure 3C**). The fact that the successive spikes occurred well within an action potential refractory period of individual cells (≤ 1msec) and project distinct waveforms into the neighboring channels suggest distinct cellular sources of each spike on the complex waveforms **(Figure 3C)**. In addition, spikes with smaller amplitude were sometimes evident in the template (*e.g.* a3 in **Figure 3C**). These spikes were most likely generated by distant cells. Networks varied in size, from 1 to 8 cells **(Figure 3F)**.

**Figure 3.**
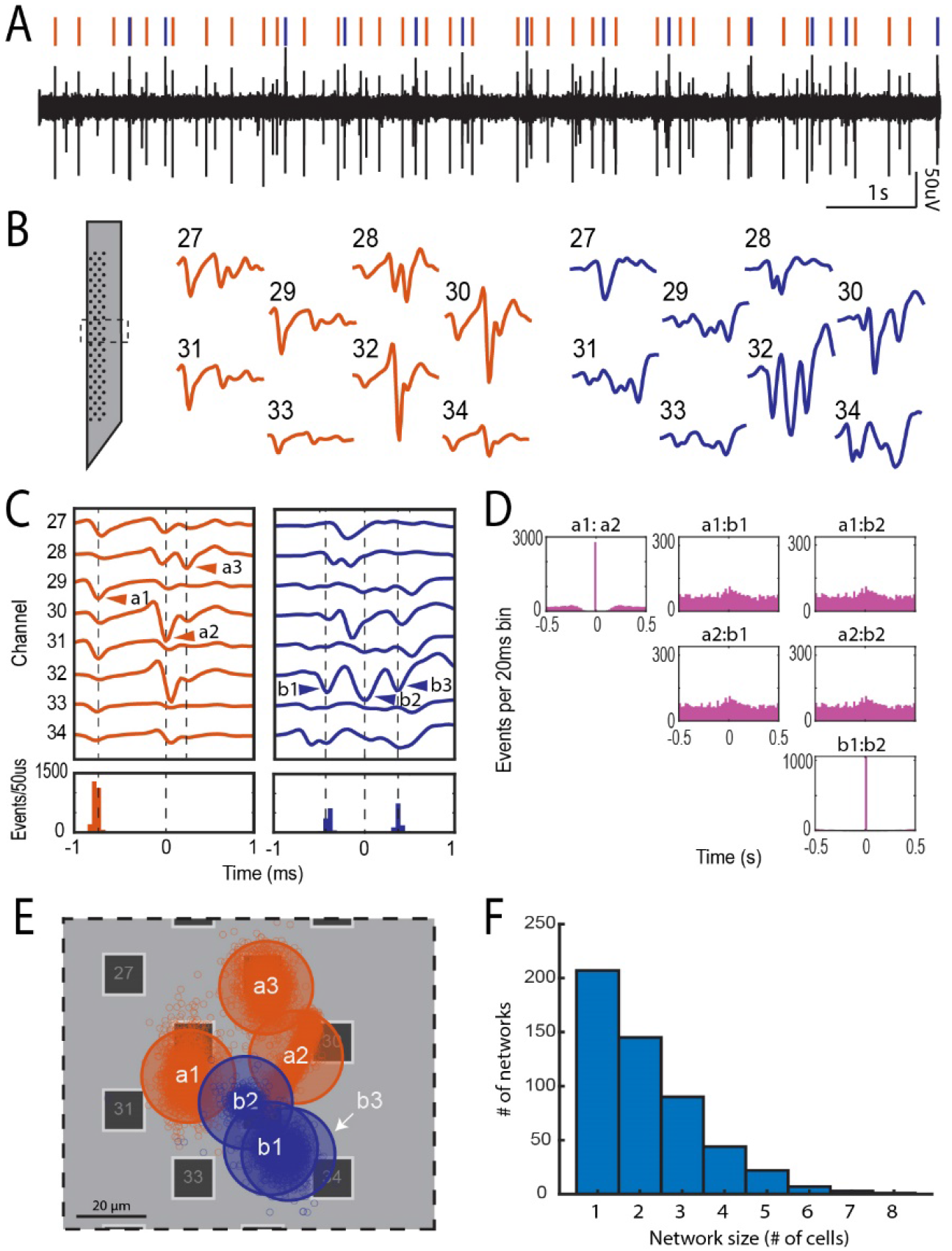
Local network activity in the adrenal medulla. A. Single channel recording showing multiple spikes (downward deflections). B. Waveforms from multiple electrodes from the experiment in panel A were sorted to derive two distinct templates shown in orange and blue. The corresponding waveforms in the single channel recording in panel A are indicated by orange and blue vertical lines. The templates displayed complex waveforms comprised of multiple spikes occurring in fixed succession. C. Some of these spikes were identified as units on their own by the sorting algorithm (e.g. a1 and a2, and b1, b2 and b3), and others were manually identified from other templates derived from another unit (e.g. a3). A temporal histogram aligned to the reference unit in the template (time = 0 ms), shows a precise firing pattern of the units (a3 was excluded from this analysis). D. A lower time resolution temporal histogram (20 ms per bin) shows a 1 to 1 correlation between a1:a2 and b1:b2, but no correlation between a and b (the total number of events for each unit are: a1=2818, a2=2875, b1=1061 and b2=1061). E. Shows the approximate location of the cells associated with each network in C. The small circles represent the theoretical locations of individual identified units. Transparent large circles represent 20 μm diameter cells centered on the centroids of the cloud of points associated with a particular unit. F. Network size distribution from a total of 40 preparations.

Units in the same template (e*.g.,* a1 and a2 or b1 and b2) displayed a high temporal correlation (>90%, a1:a2 and b1:b2 in **Figure 3D**) indicating strongly coupled components in a single network. Despite being recorded at the same recording sites, units from different templates were completely uncorrelated (a1:b1, a1:b2, a2:b1 and a2:b2 in **Figure 3D**). By estimating the two-dimensional coordinates for each unit (cell), we found that the units in the different templates could be spatially overlapping **(Figure 3E)**. Thus, despite sharing the same physical space, sets of cells displayed unique firing rates and were independent from one another, suggesting they are members of different local networks.

Network synchronization occurred over a wide time frame, from hundreds of microseconds to a few milliseconds, even in the same cellular network (**Figure 4A, D**). Correlation indices between members of a local network was usually close to 1 (**Figure 4A, bottom graph**), indicating that coupling rarely failed. There was a wide range of distances between synchronized cells (**Figure 4D)**, from as small as a cell diameter (20 μm, examples in **Figure 4B**) to as large as 500 μm (example in **Figure 4C**) . The finding that networks can be distributed over hundreds of microns indicates that they can be comprised of cells across different cell clusters.

**Figure 4.**
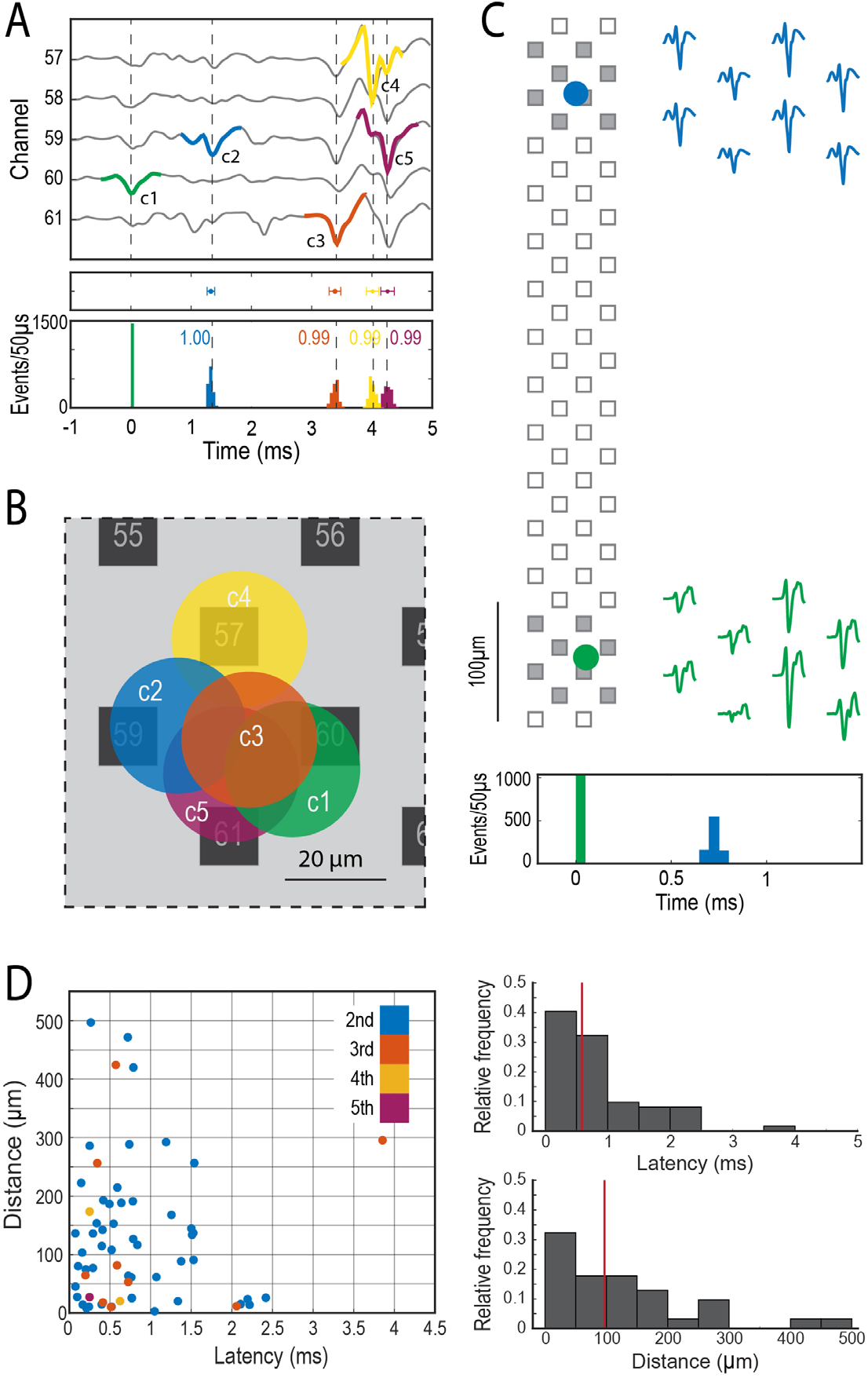
Synchronized firing in the adrenal medulla. A. Firing sequence of a group of clustered cells, Top panel shows an extended template of the first unit (−1 to 5 ms, c1 in green) displaying the firing sequence of the later units (c2 to c5, blue, orange, yellow and purple), middle panel shows the mean latency ± SD (mean ± 2SD, c2=1.33±0.06 ms, c3=3.39±0.10 ms, c4=4.01±0.10 ms and c5=4.26±0.11 ms) in reference to c1, bottom panel shows the histograms in 50 μs bins of the firing distribution between the first cell and the subsequent cells in the network. B. Shows the approximate location of the synchronized cells in A, the circles represent a 20 μm in diameter cell. C. Firing synchronization between two distant cells, the circles represent the theoretical location of each cell over the probe, the filled squares correspond to the recording sites shown in the 2 ms templates on the right. The histogram at the bottom shows the firing distribution in 50 μs bins of the second cell (blue) in relationship to the first cell (green). D. Left panel shows the relationship between latency and distance in coupled cells. Top right histogram shows the relative frequency distribution of the latency between consecutive cell pairs (n=62, the color code matches previous panels), bottom right histogram shows the relative frequency of the distance between cell pairs. The red vertical lines mark the median, 0.59 ms for latency and 97 μm for distance.

#### Evoked activity in the adrenal medulla

To further investigate electrical parameters of these chromaffin cell networks, we electrically stimulated the splanchnic nerve with different intensities (0 to 4500 μA in 500 μA steps). We found that splanchnic nerve activation reliably evoked action potentials from chromaffin cells throughout an applied frequency spectrum of 1 Hz to 40 Hz (**Figure 5A**). Application of TTX locally to the nerve reversibly inhibited evoked activity (**Figure 5B**), thus again confirming the necessity of nerve excitability to drive chromaffin cell electrical firing. Notably, splanchnic nerve stimulation resulted in the same complex waveforms as were recorded spontaneously at the same electrodes during the basal period (solid vs dotted lines in **Figure 5C**). This demonstrates that the same cellular network is activated for evoked and basal nerve activity. The evoked units fired at fixed latencies after splanchnic nerve stimulation that ranged from 5 to 30 ms **(Figure 5D)**. The firing probability increased with higher stimulus intensities suggesting distinct thresholds **(Figure 5E)**.

**Figure 5.**
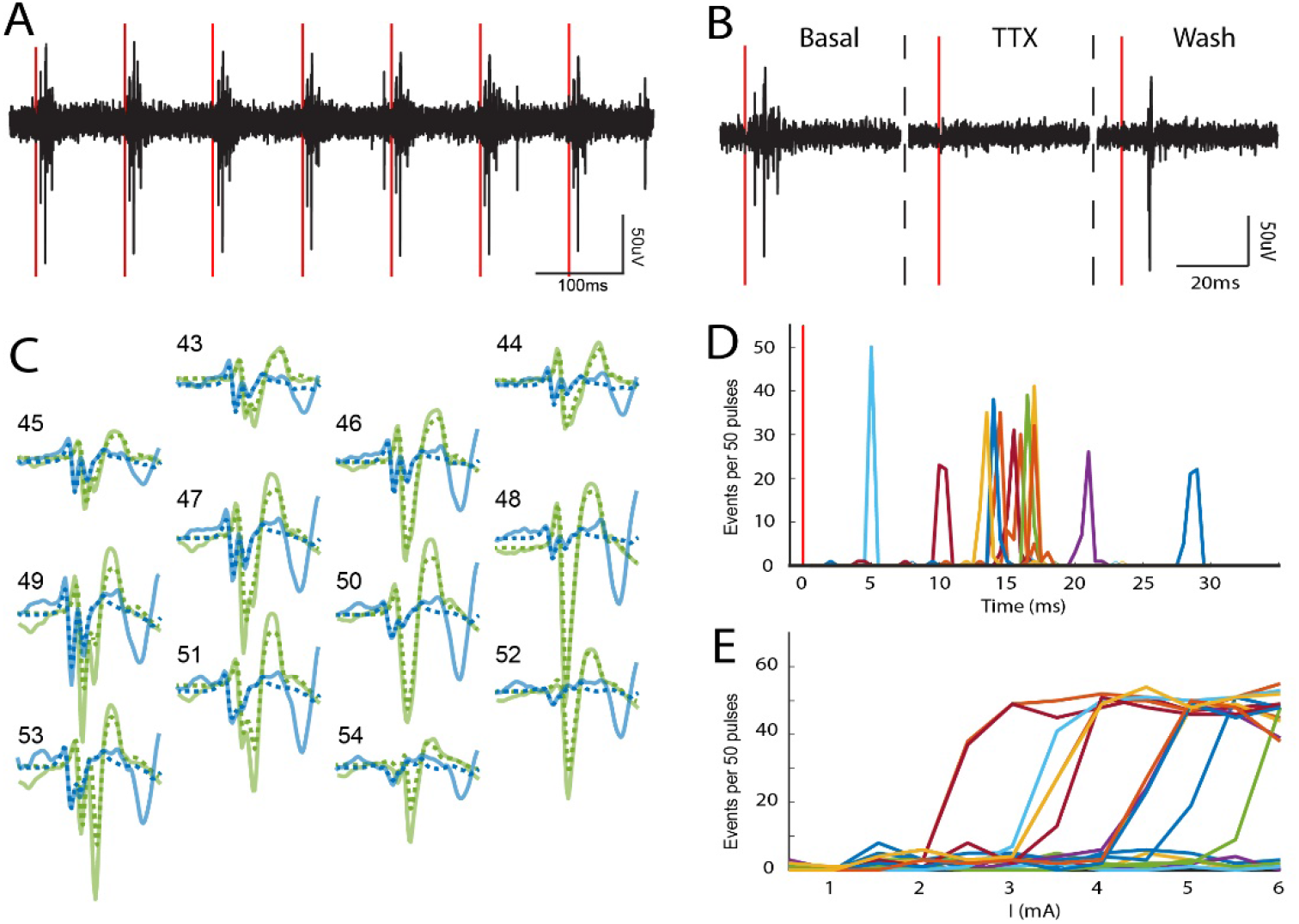
Electrically evoked chromaffin cell activity. A. Electrical stimulation of the splanchnic nerve evokes activity in the adrenal medulla. Representative recording of a single phase 10Hz-3.5mA stimulation train (red vertical lines). B. The local application of TTX to the splanchnic nerve reversibly blocked the evoked action potentials in the adrenal medulla. C. Shows the evoked templates (solid lines) from two units, these templates are identical to their spontaneous counterparts (dotted lines). D. The evoked units present distinct latencies that range from 5 to 30 ms after the stimulus (red vertical line). E. Distinct units have different stimulus thresholds. The stimulus intensity was gradually increased from 0 to 6 mA in 0.5 mA steps.

#### Physiological stress response to insulin-induced hypoglycemia

Our ability to record chromaffin cell activity in living animals allowed us to assess the physiological response of the adrenal medulla to a stressor. Anesthetized rats were challenged with increasing doses of insulin (10, 100 and 1000 mg/kg of i.p. insulin application) to initiate a hypoglycemic stress response. The challenge provoked a graded decrease in the blood glucose (red line in **Figure 6A**) and a sympathetic nervous system response that induced activation of the splanchnic nerve and stimulation of the adrenal medulla. Correlated with the fall in blood glucose was a significant increase in the activity of a subset of chromaffin cells (**Figure 6A and 6B**), these cells displayed a 1.5× increase to their basal firing rate (95% confidence between 1.3 and 1.8 with r^2^=0.84 and *p*=0.003, **Figure 6C**), suggesting a coordinated and uniform activation of a chromaffin cell population under glucose level sensitive neural control. Activation status was not determined by spatial localization since networks that did and did not respond to hypoglycemia sometimes mapped to overlapping sites (black vs red circles in **Figure 6E**). Occasionally, waveforms changed during the hypoglycemic challenge, displaying additional voltage deflections or spikes (**Figure 6D**). The changes may reflect the recruitment of neighboring cells to the network through enhanced neuronal coupling or increased gap junction coupling (see Discussion). These changes did not occur in waveforms whose frequency was unchanged during hypoglycemia.

**Figure 6.**
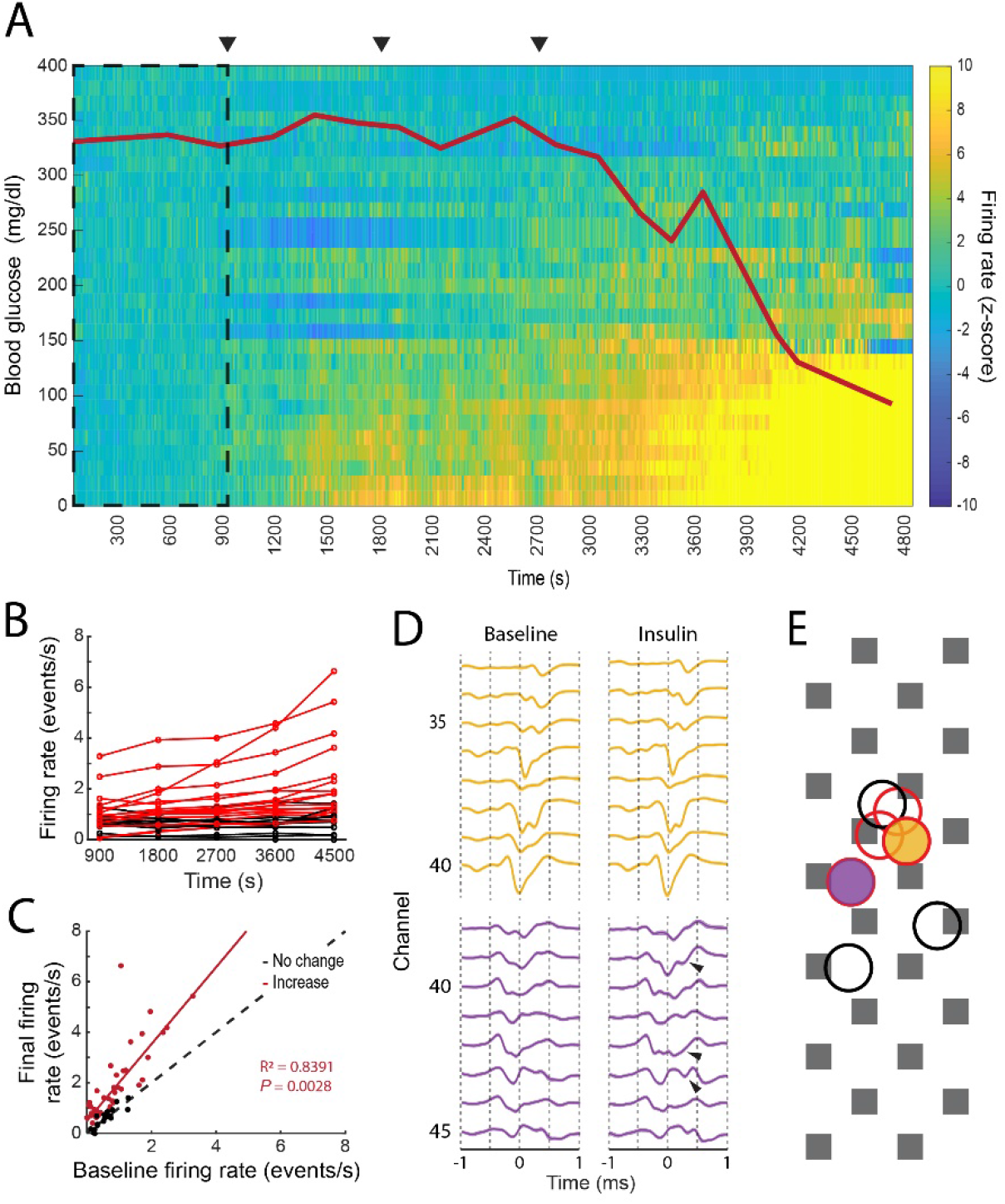
Physiological stress response in the intact adrenal medulla. A. The intraperitoneal administration of 3 increasing doses of insulin, 10, 100 and 1000 μg/kg (arrowheads), induced a decrease in the blood glucose levels (red line) that correlated with a significant increase in the firing frequency of a subpopulation of chromaffin cells. The z-score was calculated per unit in 10s bins by taking the first 900s as baseline (dashed square). The experiment was divided into five 15 minutes periods: baseline (1-900s), 10 μg/kg (901-1800s), 100 μg/kg (1801-2700s), 1000 μg/kg (2701-3600s) and final (3601-4500s) B. Individual averaged firing rate for the different periods (red indicates units whose final averaged firing rate increased by more than 2 times its baseline’s SD or averaged z-score above 2, black indicates the units that presented no change between its baseline firing rate and final firing rate or an averaged z-score between ±2). C. Linear regression between the baseline firing rate vs final firing rate for 64 units recorded from 3 preparations. The coefficients for the linear fitting are 1.523 (1.265, 1.78 with 95% confidence bounds), the 45 degree dashed line represent no change between the two evaluated periods. D. Waveform changes during the low glucose period. Top template (yellow) displayed no changes between the baseline and the 1000 μg/kg periods, whilst additional spikes (arrows) were found in the bottom unit in comparison to its baseline template, both units presented a statistically significant increase in their firing rate between the two periods. E. Firing rate change does not correlate with the spatial localization of the cells. The circles represent the theoretical localization of the cell bodies of 20μm in diameter chromaffin cells. Red and black circles represent cells whose firing rate increased or decreased respectively, yellow and purple filled circles indicate the localization of the units shown in D.

In contrast to the varied response to hypoglycemia, euthanasia induced by an overdose of general anesthetic at the end of the experiment was preceded by a generalized increase of electrical activity in virtually all recorded units (cells) of the adrenal medulla (**Figure 7A**) until activity abruptly halted. Some cells attained firing rates greater than 20 Hz for several seconds. In contrast to the stress response evoked by low glucose levels, there was no correlation between the initial and final firing rate (**Figure 7B**). Denervation almost completely prevented euthanasia-associated increase in firing (**Figure 7C**). These data demonstrate that that the increase in unit firing upon euthanasia is driven by neuronal input from the splanchnic nerve, and not by autonomous chromaffin cell activation.

**Figure 7.**
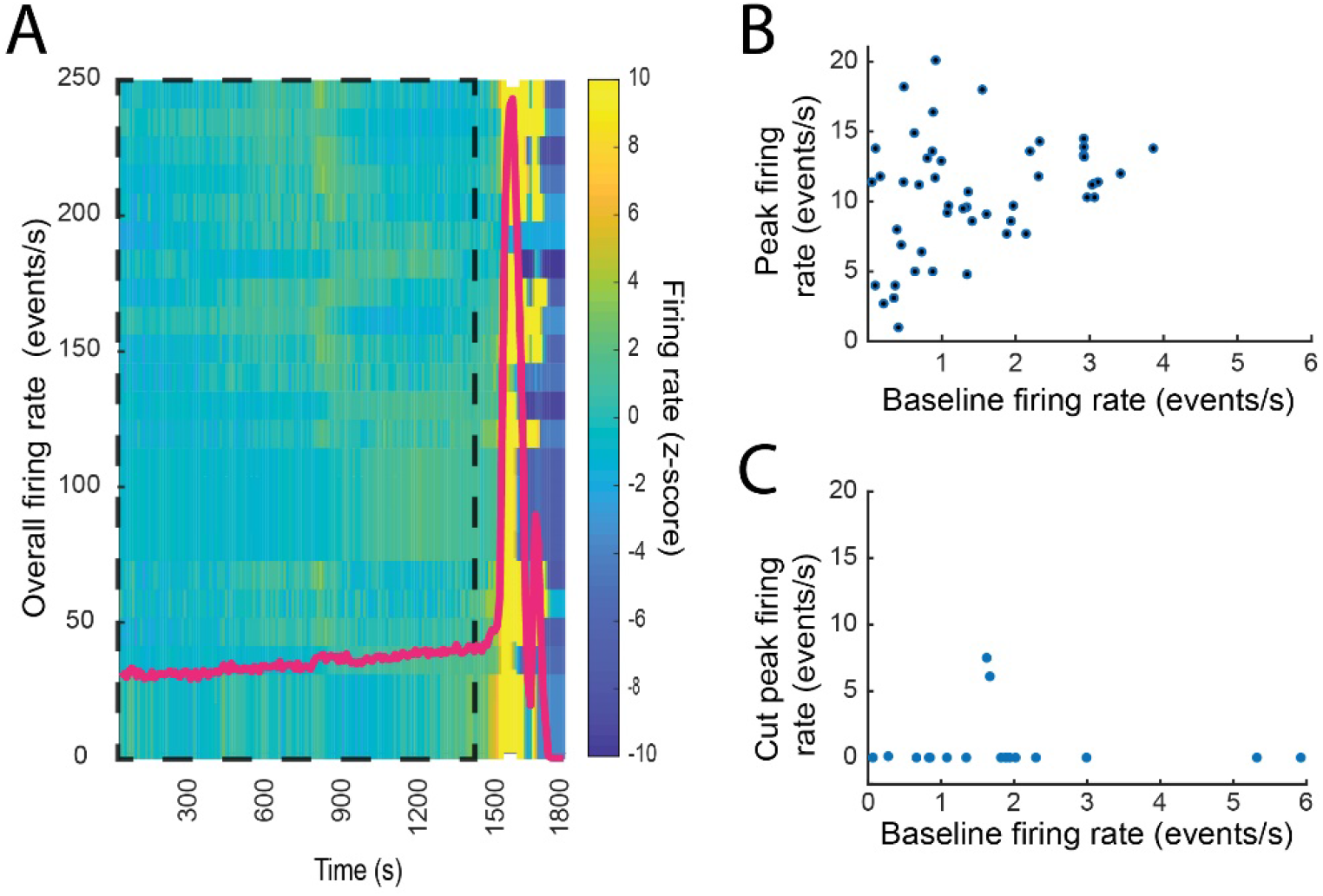
Physiological stress response to respiratory arrest in the intact adrenal medulla. A. Generalized increase in the firing rate of the chromaffin cells was observed at the moment of respiratory arrest due to anesthesia overdose. B. Relation between the baseline firing rate and the peak firing rate after respiratory arrest. C. Denervation abolishes the generalized firing rate increase after respiratory arrest.

### Discussion

We have adapted extracellular multi-electrode array technology, which had been developed to probe neuronal networks in the central nervous system, to detect adrenal chromaffin cell activity in living anesthetized rats. This approach enabled us to accurately identify and track the electrical activity of individual chromaffin cells in the living anesthetized animal. To our knowledge, this is the first application of the technology to an endocrine tissue. As discussed below, the methodology was successful in recording from as many as 40 cells simultaneously for tens of minutes in animals under basal conditions and during stimulation of the splanchnic nerve. The study revealed independent networks of chromaffin cells that had varied responses to stress, suggesting that chromaffin cells *in vivo*, rather than acting homogenously, have nuanced responses to central nervous system activation.

#### Chromaffin cell activity in living, anesthetized rats is entirely driven by splanchnic nerve stimulation

The frequency of spontaneous action potentials of individual cells varied ~0.2 to 4 Hz with a median of 1.36 Hz (**Fig. 2**). Activity was entirely driven by the splanchnic nerve. Either severing the nerve or applying tetrodotoxin directly to the nerve abolished chromaffin cell spontaneous activity. Furthermore, splanchnic nerve stimulation evoked templates of waveforms (see below) virtually identical to those observed during spontaneous activity. There was no evidence for intrinsic electrical pacemaker activity. These findings are consequential. Since secretion is almost entirely dependent upon elevation of cytosolic Ca^2+^ upon depolarization, it indicates that adrenal medullary output is tightly regulated by the nervous system rather than being autonomous or driven by acute hormonal factors.

#### Multicellular chromaffin cell networks

The multielectrode arrays enabled the generation of a template of electrode responses to the action potential of a cell with temporally associated wave forms (Pachitariu, Steinmetz et al. 2016). The method had extraordinary temporal resolution with 30,000 measurements/sec. Coupled responses separated by less than 100 μs to a few milliseconds were readily detected (**Fig. 3**). The analysis required amassing thousands of events over time. These waveforms contained highly temporally correlated contributions from other cells. The analysis revealed networks as large as 8 cells (**Fig. 3**) that were stable for tens of minutes. The network sizes were likely underestimated because of the two-dimensional layout of the electrodes.

What accounts for the coupling? One possibility is electrical coupling through gap junctions. Gap junctions have been well documented in adrenal medullary slices (Moser 1998, Martin, Mathieu et al. 2001, Hill, Lee et al. 2012, Guerineau 2018) and *in vivo* preparations (Grynszpan-Winograd 1982, Desarmenien, Jourdan et al. 2013). Gap junctions complement synaptic transmission to increase catecholamine secretion (Guerineau 2018). However, gap junction coupling likely accounts for only a small fraction of the action potential-coupled cells detected in the present study. Gap junctions of adrenal chromaffin cells are generally high resistance, and unable to conduct sufficient current between cells to transmit action potentials from one cell to another (coupling ratios significantly less than one) (Moser 1998, Martin, Mathieu et al. 2001). Furthermore, the latency of transmission of Ca^2+^ waves between coupled cells is 40 ms (Martin, Mathieu et al. 2001), much too slow to account for action potential latencies in the waveforms of milliseconds or less as detected in the present study. Although, gap junctions may be important for metabolic coupling and fine tuning of the secretory responses, they are not the primary mechanism for action potential-coupling.

It is much more likely that the low latency coupling in a network is neuronal as has been previously suggested (Kajiwara, Sand et al. 1997). Networks may reflect innervation of multiple cells by a single nerve fiber. Indeed, unmyelinated axons emerge from Schwann cell envelopment to innervate individual chromaffin cells. Neuronal coupling is suggested by electron microscope studies that show nerve terminal branching (Grynszpan-Winograd 1974) and nerve endings making synapses with more than one cell (Coupland 1965). Light microscopy demonstrates nerve fibers with at least one and sometimes several boutons per chromaffin cell (Kajiwara, Sand et al. 1997). The median distance and median latency between coupled cells were 95 μm and 0.59 ms, respectively (**Figure 4D**). By combining these two measurements, one estimates a conduction speed of 0.16 m/s, which is appropriate for small, unmyelinated axons (Hoffmeister, Janig et al. 1991, Iijima, Matsumoto et al. 1992, Chereau, Saraceno et al. 2017). The fixed firing sequence in a network also supports the notion of propagation along nerve fibers and indicates activation of postsynaptic chromaffin cells in a specific order. Coupling between neighboring cells sometimes had latencies of several milliseconds (**Figure 4D, left panel**), longer than would be expected by single fiber neuronal coupling. These longer latencies could reflect network innervation by two or more axons whose activities are coupled in the CNS.

It is attractive to suggest that coupled cells are defined by a cell cluster, the anatomical unit in the adrenal medulla (Hillarp 1946, Kajiwara, Sand et al. 1997). Our findings suggest that this is often not the case. The diameter of histological cell clusters is approximated 100 μm (**Figure 1B,C**). Forty-seven percent of coupled cells were within 100 μm of each other (**Figure 4D**); these pairs could reside in the same cluster. However, a significant fraction of synchronized cells (18%) were separated by greater than 200 μm and there were four cell pairs (6%) separated by greater than 400 μm. Furthermore, anatomical cell clusters likely do not limit coupling to one network since independent networks sometimes overlapped spatially (**Fig. 3**). Thus, a chromaffin cell network is not defined by a cell cluster, but rather by the temporal synchronization of a group of chromaffin cells regardless of their localization.

#### Nuanced response to physiological stress

The *in vivo* recordings allowed for the first time the simultaneous analysis of many individual cell responses to physiological stress. Rats were subjected to insulin shock-induced hypoglycemia. As expected, there was an increase in chromaffin cell firing rate coincident with the hypoglycemia as part of the stress response that seeks to maintain blood glucose. Surprisingly, the increase did not occur uniformly (**Fig. 6**). In some networks the increase was vigorous and sustained but in others it was short-lived or did not occur. In fact, networks that did and did not respond were sometimes spatially overlapping. These results suggest that there is a significant degree of information processing, probably in the central nervous system, that is responsible for the nuanced responses. Indeed, recent neuron tracing experiments indicate that there are extensive neuronal circuits originating in the brain stem and cerebral cortex whose final outputs are the adrenal medulla via the splanchnic nerve (Strack, Sawyer et al. 1989, Dum, Levinthal et al. 2019). Electrophysiological studies also indicate discriminating responses emanating from the CNS (Morrison and Cao 2000). The nuanced response to metabolic stress contrasts with the response to stress induced by agonal respiratory arrest (**Fig. 7**). In this case, there was a uniform increase in chromaffin cell firing rate followed by cessation, presumably because of central neuronal activation at onset of loss of CNS function. The notion that the stress response is varied and dependent upon the stimulus is consistent with previous analyses (Goldstein and Kopin 2008).

The adrenal medulla and islets are both endocrine tissues. However, there are striking differences in the functional organization of chromaffin cells in the adrenal medulla and insulin-secreting β-cells in pancreatic islets. The main signal for secretion in the adrenal medulla is neuronal through splanchnic nerve preganglionic cholinergic innervation that activates nicotinic ionic channels on chromaffin cells. The present study demonstrates that chromaffin cells are organized in small cellular networks that are coupled mainly through neuronal innervation with latencies between cells of less than a millisecond to several milliseconds. In contrast, the main stimulus for insulin secretion is the extracellular glucose concentration, which indirectly increases cytosolic Ca^2+^. β-Cell responses are coupled through the extracellular glucose concentration and gap junctions between cells with latencies of many seconds (Meissner 1976, Charollais, Gjinovci et al. 2000, Zhang, Galvanovskis et al. 2008, Bertram, Satin et al. 2018). Post-ganglionic stimulation by sympathetic and parasympathetic nerves modulates the response to glucose through relatively slow acting GTP-coupled receptors.

This initial study with a multielectrode array in the adrenal gland *in vivo* raises many questions and future research opportunities. We did not delve deeply into the post-synaptic pharmacology of synaptic transmission. However, we did investigate the role of nicotinic cholinergic receptors, the major rapid excitatory pathway in the adrenal medulla. The intraperitoneal administration of the nicotinic cholinergic antagonist, mecamylamine, inhibited transmission in approximately 50% of the cells. It is unlikely that pituitary adenylate cyclase-activating peptide (PACAP), which is present together with acetylcholine in splanchnic nerve terminals in the adrenal medulla, was alone responsible for chromaffin cell action potentials. Although its release is essential for the strong secretory response to intense stress (Przywara, Guo et al. 1996, Hill, Chan et al. 2011, Smith and Eiden 2012, Stroth, Kuri et al. 2013, Eiden, Emery et al. 2018), the cellular response to PACAP is through GTP-coupled receptor activated pathway that can take seconds to be manifest (Przywara, Guo et al. 1996) and alone does not stimulate action potentials (Hill, Chan et al. 2011). The incomplete inhibition of synaptic transmission by a nicotinic antagonist raises the possibility that the pharmacology of synaptic transmission is not uniform between chromaffin cells as has been previously suggested (Chowdhury, Guo et al. 1994).

It is possible that different cell networks have different secretory outputs. Chromaffin cells contain and secrete epinephrine (the major catecholamine in the adrenal medulla) or norepinephrine depending the cellular expression of the epinephrine synthesizing enzyme, phenylethanolamine-N-methyl transferase. There is evidence that different inputs from the CNS selectively cause epinephrine or norepinephrine secretion (Morrison and Cao 2000). Secretory granules (chromaffin granules) in chromaffin cells contain not only catecholamine but many peptides and hormones (for review (Winkler, Apps et al. 1986)) including enkephalins (Viveros, Diliberto et al. 1979), NPY, chromogranins, tissue plasminogen factor (tPA) (Parmer, Mahata et al. 1997) and plasminogen activator inhibitor-1 (PAI-1) (Jiang, Gingles et al. 2011). The mix of granule proteins differs from cell to cell (Patzak, Bock et al. 1984, Weiss, Bittner et al. 2014, Abbineni, Bittner et al. 2018). It is possible that one of the purposes of the nuanced network responses is to modulate the cocktail of substances released into the blood during stress.

### Materials and Methods

All animal handling and experimental procedures associated with or performed in this study followed National Institutes of Health (NIH) animal use guidelines and were approved by the Institutional Animal Care & Use Committee (IACUC) at University of Michigan (Approval Number PRO00009446). In addition, all research performed in this study complied with the Institutional Biosafety Committee at University of Michigan (Approval Number IBCA 00000592).

#### Animals and surgery

All the experiments were carried out in female and male Sprague Dawley rats between 90 and 180 days old, housed in an enriched environment in a 12 hr light–12 hr dark cycle and climate-controlled room (22°C) with free access to water and food. On the day of experimentation, subjects were transferred to an anesthetic induction chamber and the isoflurane anesthetic applied via a low flow vaporizer (SomnoFlo, Kent Scientific). Anesthesia was confirmed repeatedly during experiments by testing absence of a foot pain reflex. Surgery to reveal the adrenal gland was performed by making an incision through the left abdominal wall. Excess adipose tissue surrounding the adrenal gland was carefully removed. The adrenal gland was then immobilized to avoid recording artefacts associated with breathing or tissue movements and for prolonged stability of electrode placement by gently raising the adrenal and fixing placement with ring tipped forceps attached to a micromanipulator. Electrical stimulation of the intact splanchnic nerve was performed using a bipolar platinum hook electrode placed around the nerve. Multi-site silicon electrodes (64 recording site M1 probe, Cambridge NeuroTech) were introduced under hydraulic micromanipulator control (Narashige) through a small incision made into the gland’s external capsule, which allowed stable recordings of multi-site electrical activity for extended periods (**Figure 1C**).

#### Electrophysiological recordings

All the experiments were carried out with the acute version of Cambridge Neurotech’s M1 probe (**Figure 1C**), paired with two 32 channel Intan RHD2132 headstages connected to the Open Ephys hardware and software (Siegle, Lopez et al. 2017). Prior to each experiment, the electrode impedance to a 1KHz sine wave was measured and recording sites with impedance > 200KΩ were excluded from the analysis. During the experiments, the 64 channels were simultaneously recorded at 30 ksamples/s and filtered from 300 to 6000 Hz with a digital 2nd-order Butterworth filter. Finally, the signal was common average referenced and stored for offline analysis.

#### Electrical stimulation

Pulse trains of 10 single phase 100 μs constant current pulses at 1-40 Hz were generated by triggering a current stimulus isolation unit with a TTL signal from a PulsePal Gen2 (Sanworks) controlled through the Open Ephys graphical user interface (GUI). Current intensity was increased from 0.0 to 4.5 mA in 0.5 mA steps following delivery of 5 pulse trains at each intensity.

#### Spike sorting and analysis

Spike sorting was performed offline using Kilosort2 scripts and manually cured in Phy (Pachitariu, Steinmetz et al. 2016). A unit was classified as “good” when it displayed a constant firing rate during the basal period and few to no refractory period violations. After curing, homemade scripts were used to process and analyze spike’s timestamps and IDs. A unit’s template was obtained by extracting and averaging a ±3ms time window for every spike that corresponds to that specific unit from each channel in the original recording (waveform). The firing rate was calculated by counting the number of events a unit had in 10 s bins. The correlation index between units was calculated by dividing the number of times a unit fired within ±10 ms of the occurrence of the reference unit. This was analyzed throughout the total number of spikes the reference unit presented during an analysis period.

#### Pharmacology

The voltage-gated sodium channel antagonist tetrodotoxin (TTX) was locally applied by soaking a small cotton ball into a 10μM TTX solution and placing it onto the exposed splanchnic nerve. The nicotinic receptor antagonist mecamylamine (5 mg/kg, Sigma-Aldrich: M9020) was systemically administered with an intraperitoneal injection. Hypoglycemia was induced with three increasing intraperitoneal human insulin doses spaced by 15 minutes (10, 100 and 1000 μg/kg, Sigma-Aldrich: I9278), during this procedure blood glucose was periodically measured with an Accu-Chek meter (Roche) using blood samples obtained from the tail vein.

## Acknowledgments

This work was supported by the Endowment for the Basic Sciences Innovation Initiative of the University of Michigan Medical School to ELS and RWH, NIH Grant R01-170553 to RWH and NIH-NINDS Grant R01-097498 to ES. We thank Dr. Leslie Satin (/Department of Pharmacology, University of Michigan) for many illuminating discussions.

## References

Abbineni, P. S., M. A. Bittner, D. Axelrod and R. W. Holz (2018). “Chromogranin A, the major lumenal protein in chromaffin granules, controls fusion pore expansion.” J Gen Physiol 151(2): 118–130.

Bertram, R., L. S. Satin and A. S. Sherman (2018). “Closing in on the Mechanisms of Pulsatile Insulin Secretion.” Diabetes 67(3): 351–359.

Charollais, A., A. Gjinovci, J. Huarte, J. Bauquis, A. Nadal, F. Martin, E. Andreu, J. V. Sanchez-Andres, A. Calabrese, D. Bosco, B. Soria, C. B. Wollheim, P. L. Herrera and P. Meda (2000). “Junctional communication of pancreatic beta cells contributes to the control of insulin secretion and glucose tolerance.” J Clin Invest 106(2): 235–243.

Chereau, R., G. E. Saraceno, J. Angibaud, D. Cattaert and U. V. Nagerl (2017). “Superresolution imaging reveals activity-dependent plasticity of axon morphology linked to changes in action potential conduction velocity.” Proc Natl Acad Sci U S A 114(6): 1401–1406.

Chow, R. H., L. von Ruden and E. Neher (1992). “Delay in vesicle fusion revealed by electrochemical monitoring of single secretory events in adrenal chromaffin cells.” Nature 356(6364): 60–63.

Chowdhury, P. S., X. Guo, T. D. Wakade, D. A. Przywara and A. R. Wakade (1994). “Exocytosis from a single rat chromaffin cell by cholinergic and peptidergic neurotransmitters.” Neuroscience 59(1): 1–5.

Colomer, C., A. O. Martin, M. G. Desarmenien and N. C. Guerineau (2012). “Gap junction-mediated intercellular communication in the adrenal medulla: an additional ingredient of stimulus-secretion coupling regulation.” Biochim Biophys Acta 1818(8): 1937–1951.

Colomer, C., L. A. Olivos Ore, N. Coutry, M. N. Mathieu, S. Arthaud, P. Fontanaud, I. Iankova, F. Macari, E. Thouennon, L. Yon, Y. Anouar and N. C. Guerineau (2008). “Functional remodeling of gap junction-mediated electrical communication between adrenal chromaffin cells in stressed rats.” J Neurosci 28(26): 6616–6626.

Coupland, R. E. (1965). “Electron microscopic observations on the structure of the rat adrenal medulla: II. Normal innervation.” J Anat 99(Pt 2): 255–272.

Coupland, R. E. and R. L. Holmes (1958). “The distribution of cholinesterase in the adrenal glands of the rat, cat and rabbit.” J Physiol 141(1): 97–106.

Desarmenien, M. G., C. Jourdan, B. Toutain, E. Vessieres, S. G. Hormuzdi and N. C. Guerineau (2013). “Gap junction signalling is a stress-regulated component of adrenal neuroendocrine stimulus-secretion coupling in vivo.” Nat Commun 4: 2938.

Dum, R. P., D. J. Levinthal and P. L. Strick (2019). “The mind-body problem: Circuits that link the cerebral cortex to the adrenal medulla.” Proc Natl Acad Sci U S A.

Eiden, L. E., A. C. Emery, L. Zhang and C. B. Smith (2018). “PACAP signaling in stress: insights from the chromaffin cell.” Pflugers Arch 470(1): 79–88.

Gold, C., D. A. Henze, C. Koch and G. Buzsaki (2006). “On the origin of the extracellular action potential waveform: A modeling study.” J Neurophysiol 95(5): 3113–3128.

Goldstein, D. S. and I. J. Kopin (2008). “Adrenomedullary, adrenocortical, and sympathoneural responses to stressors: a meta-analysis.” Endocr Regul 42(4): 111–119.

Grynszpan-Winograd, O. (1974). “Adrenaline and noradrenaline cells in the adrenal medulla of the hamster: a morphological study of their innervation.” J Neurocytol 3(3): 341–361.

Grynszpan-Winograd, O. (1982). “Close relationship of mitochondria with intercellular junctions in the adrenaline cells of the mouse adrenal gland.” Experientia 38(2): 270–271.

Guerineau, N. C. (2018). “Gap junction communication between chromaffin cells: the hidden face of adrenal stimulus-secretion coupling.” Pflugers Arch 470(1): 89–96.

Hill, J., S. A. Chan, B. Kuri and C. Smith (2011). “Pituitary adenylate cyclase-activating peptide (PACAP) recruits low voltage-activated T-type calcium influx under acute sympathetic stimulation in mouse adrenal chromaffin cells.” J Biol Chem 286(49): 42459–42469.

Hill, J., S. K. Lee, P. Samasilp and C. Smith (2012). “Pituitary adenylate cyclase-activating peptide enhances electrical coupling in the mouse adrenal medulla.” Am J Physiol Cell Physiol 303(3): C257–266.

Hillarp, N. A. (1946). “Functional organization of the peripheral autonomic innervatiuon.” Acta Anat 4(Suppl): 1.

Hoffmeister, B., W. Janig and S. J. Lisney (1991). “A proposed relationship between circumference and conduction velocity of unmyelinated axons from normal and regenerated cat hindlimb cutaneous nerves.” Neuroscience 42(2): 603–611.

Iijima, T., G. Matsumoto and Y. Kidokoro (1992). “Synaptic activation of rat adrenal medulla examined with a large photodiode array in combination with a voltage-sensitive dye.” Neuroscience 51(1): 211–219.

Jiang, Q., N. A. Gingles, M. A. Olivier, L. A. Miles and R. J. Parmer (2011). “The anti-fibrinolytic SERPIN, plasminogen activator inhibitor 1 (PAI-1), is targeted to and released from catecholamine storage vesicles.” Blood 117(26): 7155–7163.

Kajiwara, R., O. Sand, Y. Kidokoro, M. E. Barish and T. Iijima (1997). “Functional organization of chromaffin cells and cholinergic synaptic transmission in rat adrenal medulla.” Jpn J Physiol 47(5): 449–464.

Martin, A. O., M. N. Mathieu, C. Chevillard and N. C. Guerineau (2001). “Gap junctions mediate electrical signaling and ensuing cytosolic Ca2+ increases between chromaffin cells in adrenal slices: A role in catecholamine release.” J Neurosci 21(15): 5397–5405.

Meissner, H. P. (1976). “Electrophysiological evidence for coupling between beta cells of pancreatic islets.” Nature 262(5568): 502–504.

Morrison, S. F. and W. H. Cao (2000). “Different adrenal sympathetic preganglionic neurons regulate epinephrine and norepinephrine secretion.” Am J Physiol Regul Integr Comp Physiol 279(5): R1763–1775.

Moser, T. (1998). “Low-conductance intercellular coupling between mouse chromaffin cells in situ.” J Physiol 506 (Pt 1): 195–205.

Moser, T. and E. Neher (1997). “Estimation of mean exocytic vesicle capacitance in mouse adrenal chromaffin cells.” Proceedings of the National Academy of Sciences of the United States of America 94(13)(13): 6735–6740.

Pachitariu, M., N. Steinmetz, S. Kadir, M. Carandini and H. Kenneth D. (2016). “Kilosort: realtime spike-sorting for extracellular electrophysiology with hundreds of channels.” bioRxiv: 061481.

Parmer, R. J., M. Mahata, S. Mahata, M. T. Sebald, D. T. O’Connor and L. A. Miles (1997). “Tissue plasminogen activator (t-PA) is targeted to the regulated secretory pathway. Catecholamine storage vesicles as a reservoir for the rapid release of t-PA.” J Biol Chem 272(3): 1976–1982.

Patzak, A., G. Bock, R. Fischer-Colbrie, K. Schauenstein, W. Schmidt, G. Lingg and H. Winkler (1984). “Exocytotic exposure and retrieval of membrane antigens of chromaffin granules: quantitative evaluation of immunofluorescence on the surface of chromaffin cells.” J Cell Biol 98(5): 1817–1824.

Petrovic, J., P. L. Walsh, K. T. Thornley, C. E. Miller and R. M. Wightman (2010). “Real-time monitoring of chemical transmission in slices of the murine adrenal gland.” Endocrinology 151(4): 1773–1783.

Przywara, D. A., X. Guo, M. L. Angelilli, T. D. Wakade and A. R. Wakade (1996). “A non-cholinergic transmitter, pituitary adenylate cyclase-activating polypeptide, utilizes a novel mechanism to evoke catecholamine secretion in rat adrenal chromaffin cells.” J Biol Chem 271(18): 10545–10550.

Siegle, J. H., A. C. Lopez, Y. A. Patel, K. Abramov, S. Ohayon and J. Voigts (2017). “Open Ephys: an open-source, plugin-based platform for multichannel electrophysiology.” J Neural Eng 14(4): 045003.

Smith, A. D. and H. Winkler (1972). Fundamental mechanisms in the release of catecholamines. Catecholamines. H. Blaschko and E. Muscholl. Berlin, Springer-Verlag: 538–615.

Smith, C. B. and L. E. Eiden (2012). “Is PACAP the major neurotransmitter for stress transduction at the adrenomedullary synapse?” J Mol Neurosci 48(2): 403–412.

Strack, A. M., W. B. Sawyer, K. B. Platt and A. D. Loewy (1989). “CNS cell groups regulating the sympathetic outflow to adrenal gland as revealed by transneuronal cell body labeling with pseudorabies virus.” Brain Res 491(2): 274–296.

Stroth, N., B. A. Kuri, T. Mustafa, S. A. Chan, C. B. Smith and L. E. Eiden (2013). “PACAP controls adrenomedullary catecholamine secretion and expression of catecholamine biosynthetic enzymes at high splanchnic nerve firing rates characteristic of stress transduction in male mice.” Endocrinology 154(1): 330–339.

Viveros, O. H. (1976). Mechanism of secretion of catecholamines from adrenal medulla. Handbook of Physiology Endocrinology, Sec. 7, Vol. 6. H. Blaschko, G. Sayers and A. D. Smith. Washington, D.C., American Physiological Society: 389–426.

Viveros, O. H., E. J. Diliberto, E. Hazum and K. J. Chang (1979). “Opiate-like materials in the adrenal medulla: evidence for storage and secretion with catecholamines.” Mol.Pharmacol. 16: 1101–1108.

Wakade, A. R. and T. D. Wakade (1982). “Secretion of catecholamines from adrenal gland by a single electrical shock: electronic depolarization of medullary cell membrane.” Proc Natl Acad Sci U S A 79(9): 3071–3074.

Weiss, A. N., M. A. Bittner, R. W. Holz and D. Axelrod (2014). “Protein Mobility within Secretory Granules.” Biophys J 107(1): 16–25.

Wightman, R. M., J. A. Jankowski, R. T. Kennedy, K. T. Kawagoe, T. J. Schroeder, D. J. Leszczyszyn, J. A. Near, E. J. Diliberto, Jr. and O. H. Viveros (1991). “Temporally resolved catecholamine spikes correspond to single vesicle release from individual chromaffin cells.” Proc Natl Acad Sci U S A 88(23): 10754–10758.

Winkler, H., D. K. Apps and R. Fischer-Colbrie (1986). “The molecular function of adrenal chromaffin granules: established facts and unresolved topics.” Neuroscience 18(2): 261–290.

Wolf, K., G. Zarkua, S. A. Chan, A. Sridhar and C. Smith (2016). “Spatial and activity-dependent catecholamine release in rat adrenal medulla under native neuronal stimulation.” Physiol Rep 4(17).

Zhang, Q., J. Galvanovskis, F. Abdulkader, C. J. Partridge, S. O. Gopel, L. Eliasson and P. Rorsman (2008). “Cell coupling in mouse pancreatic beta-cells measured in intact islets of Langerhans.” Philos Trans A Math Phys Eng Sci 366(1880): 3503–3523.

